# Highly effective gene inactivation in tetraploid *Xenopus laevis* with low-temperature-active engineered Cas12a

**DOI:** 10.1101/2023.07.30.550813

**Authors:** Seongmin Yun, Seohyun Kim, Juyeon Hong, Seungyun Shin, Jiwon Choi, Jeongsik Shin, Subin Choi, Hyun-Shik Lee, Seung Woo Cho, Tae Joo Park, Taejoon Kwon

## Abstract

Gene knockout using the CRISPR/Cas (clustered regulatory interspaced short palindromic repeats/CRISPR-associated protein) system revolutionized reverse genetic studies in model and non-model organisms, as almost all genetic elements can be targeted with few limitations. Although the CRISPR/Cas system with SpCas9 (Cas9 derived from Streptococcus pyogenes) remains the most popular for genome editing, another CRISPR/Cas system with Cas12a (Cpf1) has expanded its application. However, Cas12a is challenging to use in some aquatic model organisms, such as *Xenopus laevis*, because of its low activity at the temperature at which X. laevis embryos are usually raised (lower than 25°C). Recently, an engineered Cas12a called Cas12a-Ultra was developed, which has improved in vivo endonuclease activity with reduced temperature dependency. Here, we evaluated the performance of these engineered Cas12a enzymes in X. laevis embryos. We first confirmed that they were more active than SpCas9 at the low temperature at which X. laevis embryos are mostly raised (20–22°C), based on in vitro digestion experiments. Then, we evaluated in vivo activities of Cas12a-Ultra by disrupting several genes in X. laevis whose phenotypic consequences are previously reported. LbCpf1 (Cas12a derived from Lachnospiraceae bacterium)-Ultra outperformed the other enzymes, producing more than 80% of embryos with severely defective phenotypes even in low-temperature conditions. In addition, duplicated copies of two paralogous lysine demethylases (kdm5b and kdm5c) were successfully disrupted, which recapitulated the previously reported phenotypes observed upon morpholino-mediated knockdown. This study demonstrated that this engineered Cas12a is valuable for gene function studies in Xenopus and other model organisms with low growth temperatures.

## Introduction

*Xenopus laevis* is a primary model organism in cell and developmental biology because of its advantageous features, such as its evolutionary conservation with humans, the ease of obtaining more than 1000 embryos, and well-established techniques for gene and tissue manipulation (Harland & Grainger 2011). However, it is challenging to perform genetic manipulation in *X. laevis* because its generation time (1–2 years) is longer than that of other vertebrate models such as mice and zebrafish, although its diploid sister species *Xenopus tropicalis* has a shorter generation time (3–6 months) (Abu-Daya et al. 2012). Additionally, because of four copies of target genes compared with two copies in diploid species, *X. laevis* with an allotetraploid genome, hinders genetic studies (Session et al. 2016). Due to these limitations, gene function studies mainly depend on specialized antisense oligonucleotide (morpholino)-based gene perturbation methods, even though their side effects are debatable (Blum et al. 2015; Gentsch et al. 2018; Paraiso et al. 2019).

CRISPR/Cas9 (clustered regulatory interspaced short palindromic repeats/CRISPR-associated protein 9) has become a popular genome manipulation tool in various model organisms, including *Xenopus*. After this method was originally established in diploid *X. tropicalis* (Blitz et al. 2013; Nakayama et al. 2013; Shigeta et al. 2016), it was quickly expanded to effectively disrupt duplicated genes in the tetraploid *X. laevis* (DeLay et al. 2018; Kato et al. 2021; Tanouchi et al. 2022; Wang et al. 2015). Furthermore, this method was developed to introduce transgenes into the genome (Nakayama et al. 2020; Shigeta et al. 2016), enabling various disease-associated alleles to be tested in this model.

Following the identification of the Cas9 enzyme from *Streptococcus pyogenes* (SpCas9), various CRISPR-Cas systems were developed through metagenomic studies of prokaryotes (Makarova et al. 2020). Among them, Cas12a, also known as Cpf1, has been spotlighted from a genome engineering perspective because it does not require a tracrRNA and can process its precursor (Fonfara et al. 2016; Zetsche et al. 2017; Zetsche et al. 2015), and has a distinct molecular mechanism compared with Cas9 (Swarts & Jinek 2018). Cas12a derived from *Acidaminococcus* (AsCpf1) or *Lachnospiraceae* (LbCpf1) bacterium is well characterized and widely used for genome engineering. However, the enzymatic activity of natural Cas12a strongly depends on the reaction temperature, limiting its usability in aquatic model organisms such as *Xenopus* and zebrafish, which require a habitat temperature lower than 30°C (Moreno-Mateos et al. 2017). Zebrafish is more tolerant of high temperatures and therefore can be used at 34°C, but neither *X. laevis* nor *X. tropicalis* embryos tolerate high temperatures. Recently, engineered variants of Cas12a were produced using a direct evolution approach (Zhang et al. 2021). With only two amino acid changes, these variants, called AsCpf1-Ultra and LbCpf1-Ultra, have significantly improved *in vivo* activities and exhibit enzymatic activity at a wide range of working temperatures. Therefore, we hypothesized that these engineered variants would also be beneficial to use in *Xenopus*.

Here, we systematically examined the activities of these engineered Cas12a enzymes (AsCpf1-Ultra and LbCpf1-Ultra) to disrupt *in vivo* gene functions in tetraploid *X. laevis*. These two enzymes showed significantly higher *in vivo* gene disruption efficiencies than SpCas9 and another engineered Cas12a (EnGen-LbCpf1). We quantitatively measured the efficiency of these enzymes by analyzing embryo images with abnormal eye pigmentation patterns and reduced head size using an image quantification method similar to that in a previous report (Tanouchi et al. 2022). Our work reported here confirmed that AsCpf1-Ultra and LbCpf1-Ultra could be an efficient alternative to SpCas9 for CRISPR applications, such as gene disruption in *Xenopus* and other model systems requiring low growth temperatures.

## Materials and Methods

### Animals

*X. laevis* embryos were obtained and maintained according to standard protocols (Sive et al. 2010). All animal experiments were approved by the Institutional Animal Care and Use Committee (IACUC) of the Ulsan National Institute of Science and Technology (Ulsan, Republic of Korea) (UNISTIACUC-22-49).

### Cas12a proteins and mRNAs

The proteins of SpCas9, AsCpf1-Ultra originating from *Acidaminococcus sp.* BV3L6, and LbCpf1-Ultra originating from the *Lachnospiraceae* bacterium ND2006 were purchased from Integrated DNA Technologies (Coralville, Iowa, United States). Another engineered Cas12a (EnGen-LbCpf1) originating from the *Lachnospiraceae* bacterium ND2006 was purchased from New England Biolabs (Ipswich, Massachusetts, United States). AsCpf1-Ultra cDNA was cloned from an AsCpf1 lentiviral vector (Addgene #84750) by incorporating two mutations (M537R and F870L) reported in a previous study (Zhang et al. 2021) via PCR mutagenesis. Then, the cDNA was cloned into the CS108 vector for *in vitro* mRNA synthesis, and its mRNA was transcribed from the cloned plasmid using an mMESSAGE mMACHINE™ SP6 Transcription Kit (Invitrogen).

### Guide RNAs

To design guide RNAs and primers, CHOPCHOP (version 3, http://chopchop.cbu.uib.no/) (Labun et al. 2019) was used with TTTN as the PAM based on version 9.2 of the *X. laevis* genome. Additionally, the guide RNAs were searched against the *X. laevis* genome (version 9.2) downloaded from the Xenbase online genome server (http://www.xenbase.org/, RRID: SCR_003280) (Fisher et al. 2023; Fortriede et al. 2020) using Exonerate (version 2.2.0) (Slater & Birney 2005) with an allowance of two-base mismatches. All CRISPR RNAs (crRNAs) and tracrRNA were purchased from Integrated DNA Technologies (Coralville, Iowa, United States) and validated by *in vitro* digestion tests. The target sequences and primer sequences of each gene are shown in Tables 1 and 2. To test the availability of Cas12a guide RNA *in vivo*, we utilized polycistronic IN4MER crRNA templates with four crRNAs together

**Table 1.**
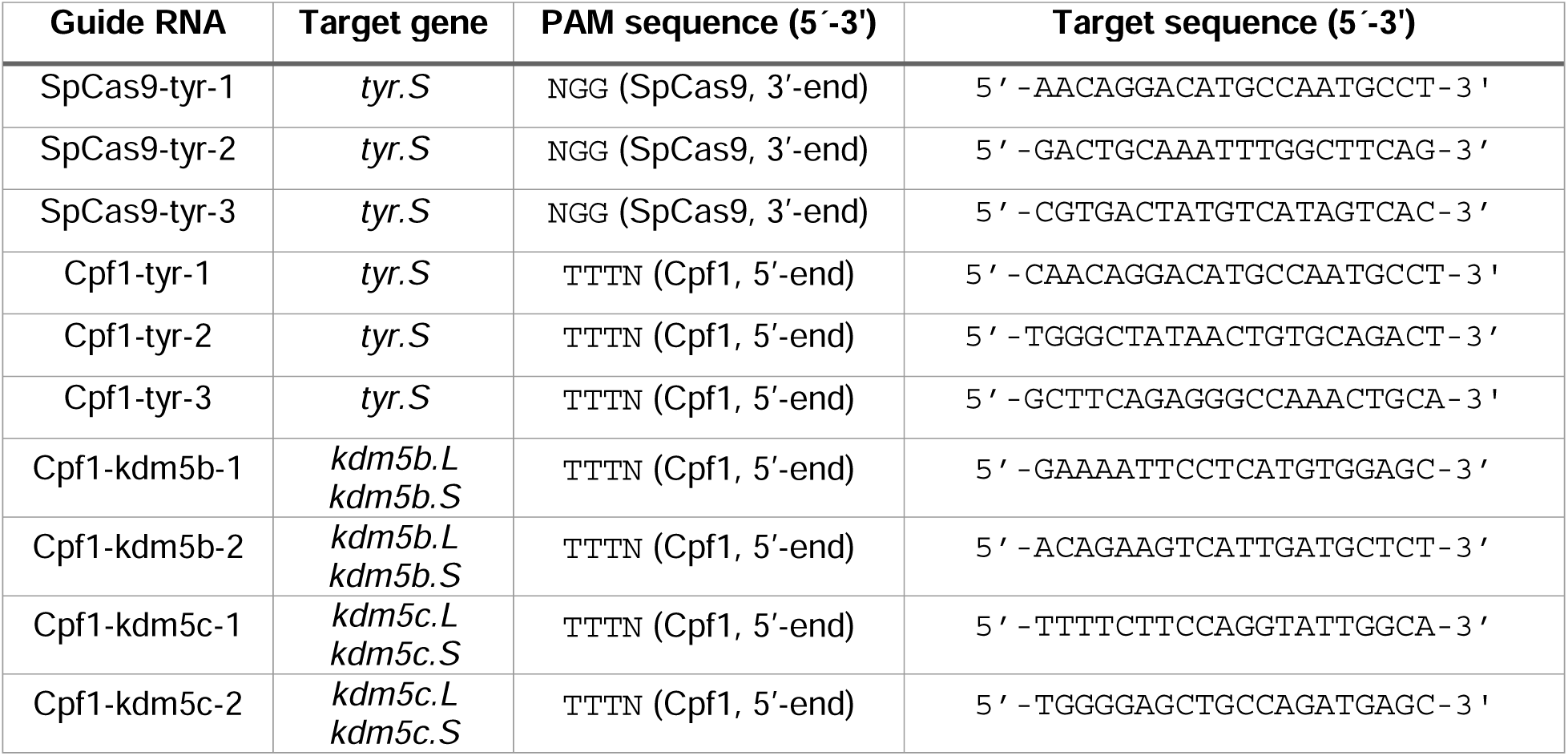
Guide RNA sequences used in this study. For *kdm5b* and *kdm5c*, guide RNAs targeting the duplicated L and S copies in the *X. laevis* genome were designed. This double targeting condition was not considered for tyrosinase because there is only one copy of the tyrosinase gene (*tyr*) in the *X. laevis* genome.

**Table 2.**
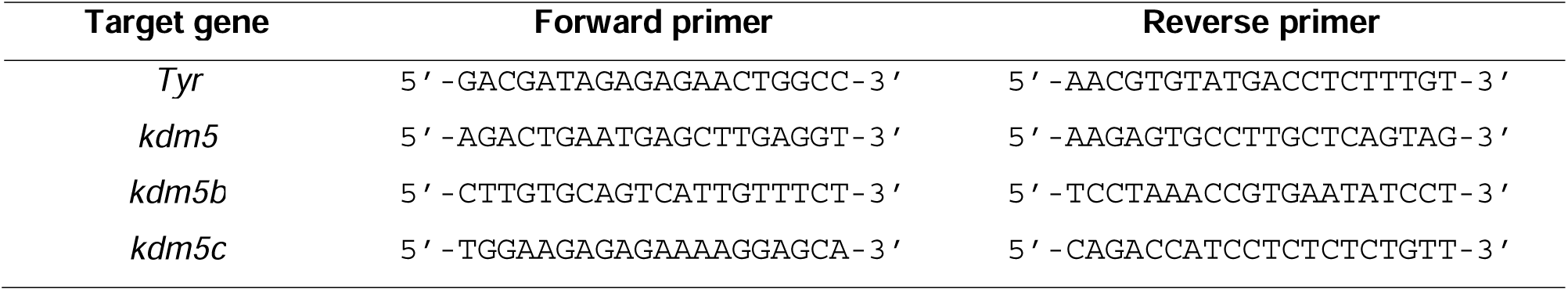
Primer sequences used in this study.

### Microinjection

Adult *X. laevis* females were injected with 400 µL of human chorionic gonadotropin (2000 IU/mL, Sigma-Aldrich) and incubated for more than 16 h to induce ovulation in an incubator at 16°C. Unfertilized eggs were collected by gently squeezing female frogs. *In vitro* fertilization was performed by stirring eggs and minced testes dissected from deeply anesthetized and euthanized male frogs in a 90 mm Petri dish. When embryos reached the 2-cell developmental stage, the jelly layer was removed using 3% cysteine solution (pH 7.8), and embryos were maintained under 1/3× Marc’s Modified Ringers (MMR). Before microinjection, 3 µM Cas protein (Cas9 and Cas12a) and 3 µM CRISPR RNAs were assembled for 10 min *in vitro* at room temperature. Ribonucleoprotein (RNP) complexes were injected into the animal-ventral region of embryos at the 2-cell stage under 1/3× MMR containing 3% Ficoll (Biosesang). We chose to inject both sides of 2-cell stage embryos to filter out embryos with abnormal development without proper early cleavages. After three hours, the injected embryos were transferred to 1/3× MMR containing gentamicin.

### Phenotype analysis of crispants embryos

Eye size and pigmentation pattern were assessed using a standardized image analysis as follows. Embryos were placed in a 90 mm Petri dish and scanned using an EPSON V850 PRO scanner at a resolution of 5400 dpi. Subsequently, the scanned images were processed using ImageJ (version 2.9.0) (Schneider et al. 2012) to crop the eyes. Specifically, the ‘color threshold’ option in ImageJ was used to filter out regions with RGB values lower than 70, thereby isolating the black pigment associated with the eyes. The number of black pixels was then measured as a proxy for eye size. The minimum eye area for a normal embryo was approximately 2000 black pixels. Consequently, eye regions exceeding this threshold were classified as ‘Normal’. Embryos exhibiting a complete absence of dark pigmentation were classified as ‘Severe’, while those in between were classified as ‘Moderate’. Head size for kdm5b and kdm5c crispants was measured manually after taking the images with a conventional microscope.

## Results

### Engineered Cas12a is fully functional at the low temperature at which *Xenopus* embryos are usually maintained

We first examined the temperature dependency of engineered Cas12a by performing an *in vitro* digestion assay (Fig. 1). Although AsCpf1-Ultra is reportedly active at 30°C (Zhang et al. 2021), this temperature is still too high to raise *X. laevis* embryos. Therefore, we tested the efficiency of *in vitro* digestion of the amplicon of *X. laevis* tyrosinase (*tyr.S*) at 18–28°C. As expected, SpCas9 had a digestion efficiency of 86–88% without temperature dependency within this range, but a relatively high concentration (3 µM) of this enzyme was required (Fig. 1A and Fig. 1B). Because one micromolar of SpCas9 is sufficient to achieve a similar efficiency at 37 °C, we speculated that the lower temperature reduces the efficiency to find the target. On the other hand, AsCpf1-Ultra (93–96%) and LbCpf1-Ultra (almost 100%) had much higher digestion efficiencies even at a concentration of 1 µM (Fig. 1C and Fig. 1D). Remarkably, the digestion efficiency of LbCpf1-Ultra was almost 100% at 20–28°C, and thus this enzyme may be more efficient in *Xenopus* embryos.

**Figure 1.**
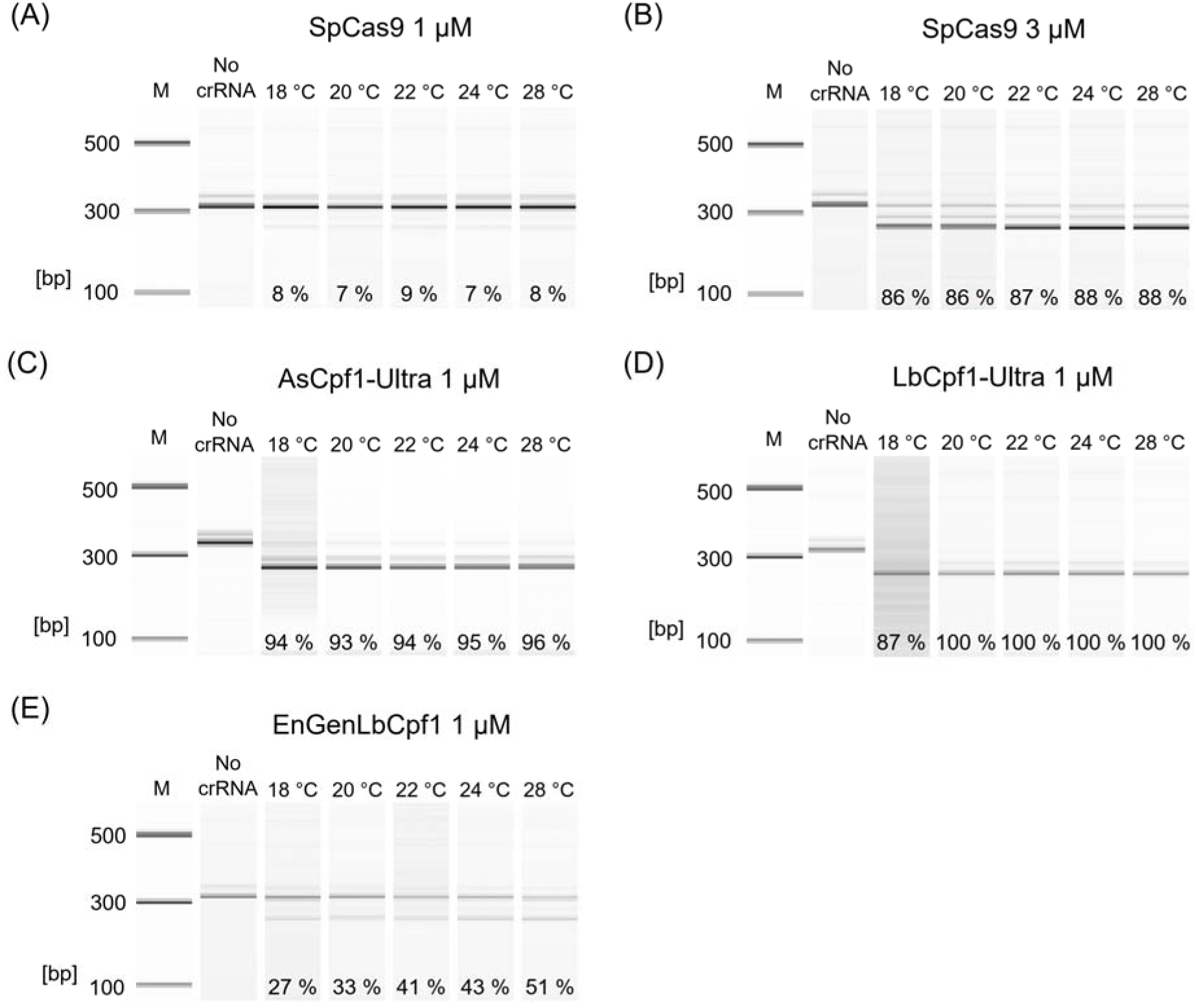
Cpf1-Ultra is functional without temperature dependency. The temperature dependency of Cpf1-Ultra was tested by assessing *in vitro* digestion of the *X. laevis* tyrosinase (*tyr*) gene (guide RNAs used in this experiment are listed in Table 1). The amount of input DNA of the reaction is measured on the lane of ‘No crRNA’ using the BioAnalyzer DNA analysis kit (Agilent Technologies, Inc.), and the residual DNA of the same size was measured to calculate the efficiency of *in vitro* cleavage, as noted at the bottom of each lane. **(A and B)** SpCas9 showed no temperature dependency in terms of its endonuclease activity, but a high concentration (3 μM) of this enzyme was required. **(C and D)** Cpf1-Ultra cut the target DNA regardless of the temperature, even at a low enzyme concentration (1 μM). **(E)** EnGen-LbCpf1, another engineered Cpf1, showed less efficient endonuclease activity than Cpf1-Ultra under the same conditions.

We also tested another engineered Cas12a enzyme from a different vendor (EnGen-LbCpf1) whose modification has not been fully disclosed. Although it exhibited digestion activity (27–51%) at the low temperatures tested, it also showed temperature dependency, with higher digestion efficiencies at higher temperatures. Therefore, we concluded that AsCpf1-Ultra and LbCpf1-Ultra are functional at the temperature at which *Xenopus* embryos are usually raised.

### Engineered Cas12a efficiently disrupts the tyrosinase gene in *X. laevis* embryos

To test the *in vivo* activity of engineered Cas12a, we formed an RNP complex comprising the Cas12a enzyme and guide RNAs targeting the *X. laevis* tyrosinase gene, and injected it into two blastomeres of the two-cell stage *X. laevis* embryos (Fig 2). Disruption of the tyrosinase gene leads to an albino-like pigmentation absence; therefore, we could measure the *in vivo* efficiency of the CRISPR-Cas system visually. To investigate the *in vivo* temperature dependency of the engineered Cas12a enzymes, one group of embryos was incubated at 20°C after injection, while another group was incubated at 28°C for one hour and then moved to 20°C. Both AsCpf1-Ultra and LbCpf1-Ultra induced severe albino-like phenotypes in both conditions, but EnGen-LbCpf1 induced mild mosaic phenotypes. We confirmed *in vivo* digestion by these enzymes based on Sanger sequencing chromatograms (Fig. 2A).

**Figure 2.**
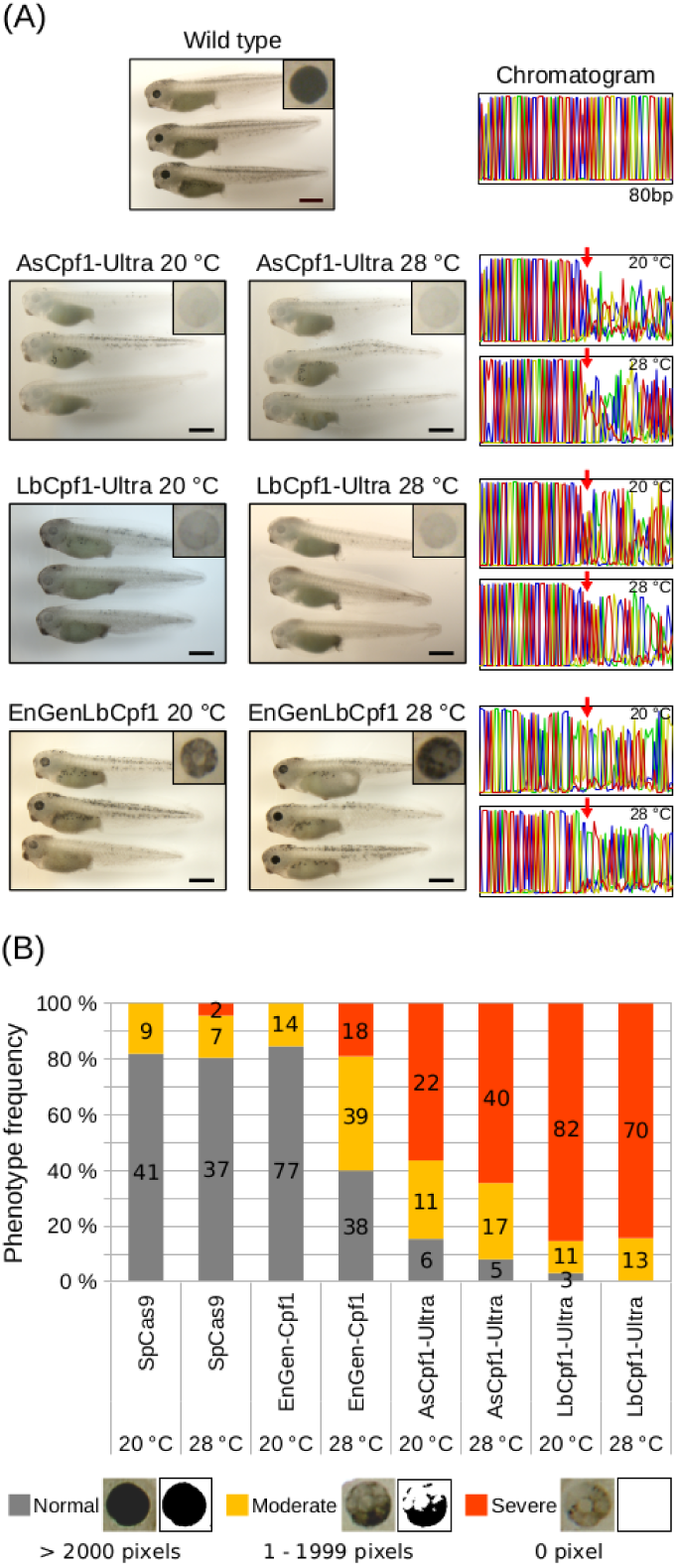
Cpf1-Ultra efficiently disrupts *X. laevis* tyrosinase gene function *in vivo*. Both 20°C (the temperature at which we typically raise *X. laevis* embryos) and 28°C (a high temperature at which *X. laevis* embryos can survive for one hour) were tested. **(A)** Wild-type tadpoles at NF stage 40 showed complete pigmentation of the eyes and skin (scale bars, 1 mm; digitally magnified eye images are also presented in the boxes), but crispants generated with Cpf1-Ultra (both AsCpf1-Ultra and LbCpf1-Ultra) showed almost no pigmentation. Crispants generated with EnGen-Cpf1 showed reduced pigmentation, but the effect was less severe than that of Cpf1-Ultra. The *in vivo* digestion of the target site (marked by a red arrow) was confirmed by Sanger sequencing. **(B)** Eye pigmentation was quantified by segmenting the eye field and counting the number of black pixels. The phenotypes were classified into three groups according to the number of dark pixels (‘Normal’ for more than 2000 pixels, ‘Moderate’ for 1–1999 pixels, and ‘Severe’ for 0 pixels). The number of examined embryos is also presented.

Although the body’s pigmentation clearly changed due to the disruption of the tyrosinase enzyme, we assessed the gene disruption efficiency more quantitatively by measuring the eye pigmentation at NF stage 40. Eye size is almost identical at a given stage; therefore, we counted the number of black pixels in segmented eye images and empirically classified them into three categories. The phenotype was designated as ‘Normal’ if the number of black pixels was the same as in control embryos (mostly more than 2,000 pixels), ‘Severe’ if no black pixels were detected, and ‘Moderate’ if there were 1–1,999 black pixels (Fig. 2B). Surprisingly, more than 80% of embryos had a ‘Severe’ phenotype and more than 96% of embryos had an abnormal phenotype using LbCpf1-Ultra, which outperformed all other enzymes tested including SpCas9 regardless to the 28°C heat shock. From this result, we concluded that LbCpf1-Ultra is highly efficient at disrupting genes in *X. laevis* even at the normal culture temperature.

### Engineered Cas12a can simultaneously disrupt duplicated genes in the tetraploid *X. laevis* genome

The *X. laevis* genome contains 8,806 homoeologous genes (two copies of genes from the L chromosome and two copies of genes from the S chromosome) (Session et al. 2016). However, the tyrosinase gene appears to be a singleton gene present only on the S chromosome; therefore, we could not validate whether engineered Cas12a can simultaneously disrupt duplicated *X. laevis* genes using this gene. Instead, we chose the lysine methyltransferase gene *kdm5c*. We previously reported that suppression of *kdm5c* expression using a morpholino reduces head size and induces an eye development abnormality (Kim et al. 2018). We also tested its singleton paralog, kdm5b, to compare functional redundancy.

We injected the LbCpf1-Ultra RNP with guide RNAs targeting *kdm5b* and *kdm5c* individually. Most *kdm5c*-disrupted embryos had small heads, as we previously observed upon knockdown using a morpholino (Fig. 3A) (Kim et al. 2018). The CRISPR-Cas system can function at various stages of development. Consequently, even if the RNP is injected at a very early stage, the mosaic phenotype becomes prevalent, hindering the application of this system for gene disruption. However, the phenotypic penetrance rate of LbCpf1-Ultra was higher than 90% (37 out of 40 in *kdm5b* crispants and 40 out of 44 in *kdm5c* crispants), which is comparable with that achieved by knockdown using a morpholino. Head size (Fig. 3B) and eye size (Fig. 3C and Fig. 3D) were significantly reduced in *kdm5b* and *kdm5c* crispants. This result proves that LbCpf1-Ultra could be an alternative option for studying gene function through disruption.

**Figure 3.**
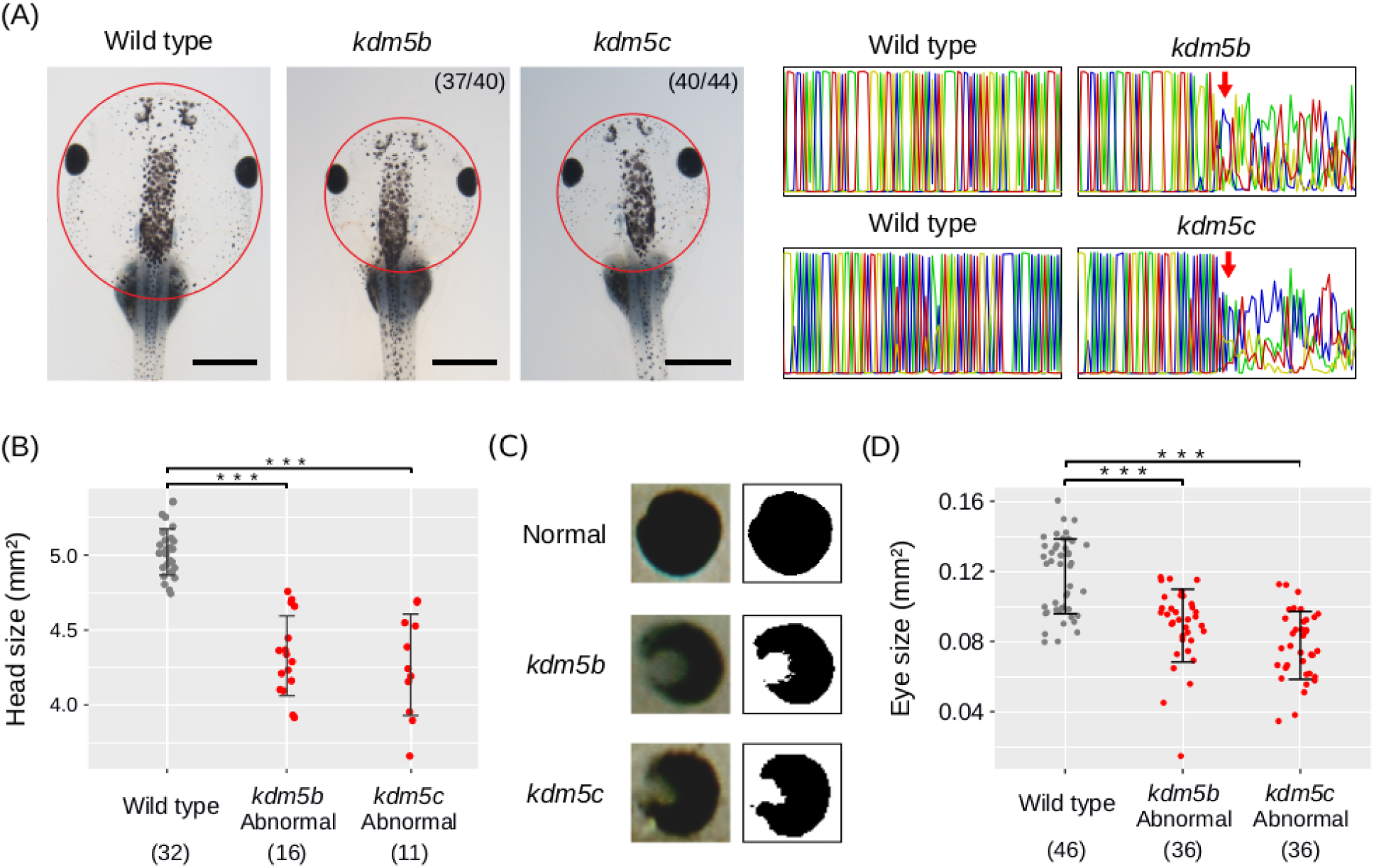
Gene disruption using LbCpf1-Ultra reproduces the phenotype observed upon morpholino-mediated knockdown. The phenotype observed upon morpholino-mediated knockdown of the histone demethylase kdm5c and its paralog kdm5b that we previously reported (Kim et al. 2018) was recapitulated. **(A)** Head size was significantly reduced in both *kdm5b* and *kdm5c* crispants. Scale bars, 1 mm. Notably, more than 90% of crispants showed consistent phenotypic outcomes. The *in vivo* digestion was confirmed by sequencing. **(B)** The head size of *kdm5*-disrupted tadpoles was significantly reduced (*** represents a p-value less than 0.001 with the two-sided Student’s t-test). **(C–D)** The eye defect was quantified by quantifying the pigmentation of the segmented eye field of NF stage 40 embryos (*** represents a p-value less than 0.001 with the two-sided Student’s t-test).

### Engineered Cas12a is not functional when injected as mRNA due to the rapid degradation of guide RNAs

Previously, it was reported that Cas12a mRNA is not functional when injected simultaneously with guide RNAs in zebrafish (Moreno-Mateos et al. 2017). To test whether this is due to low enzymatic activity, we injected the guide RNAs targeting the *X. laevis* tyrosinase gene together with AsCpf1-Ultra mRNA, which we validated with the RNP (Fig. 2), and synthetic mRNA produced via *in vitro* transcription. However, we could not observe any abnormal phenotypes (Fig. 4A). Additionally, when we checked the target site by Sanger sequencing, we found no peak disruption signature, which is usually generated when CRISPR works, as we confirmed in the RNP injection.

**Figure 4.**
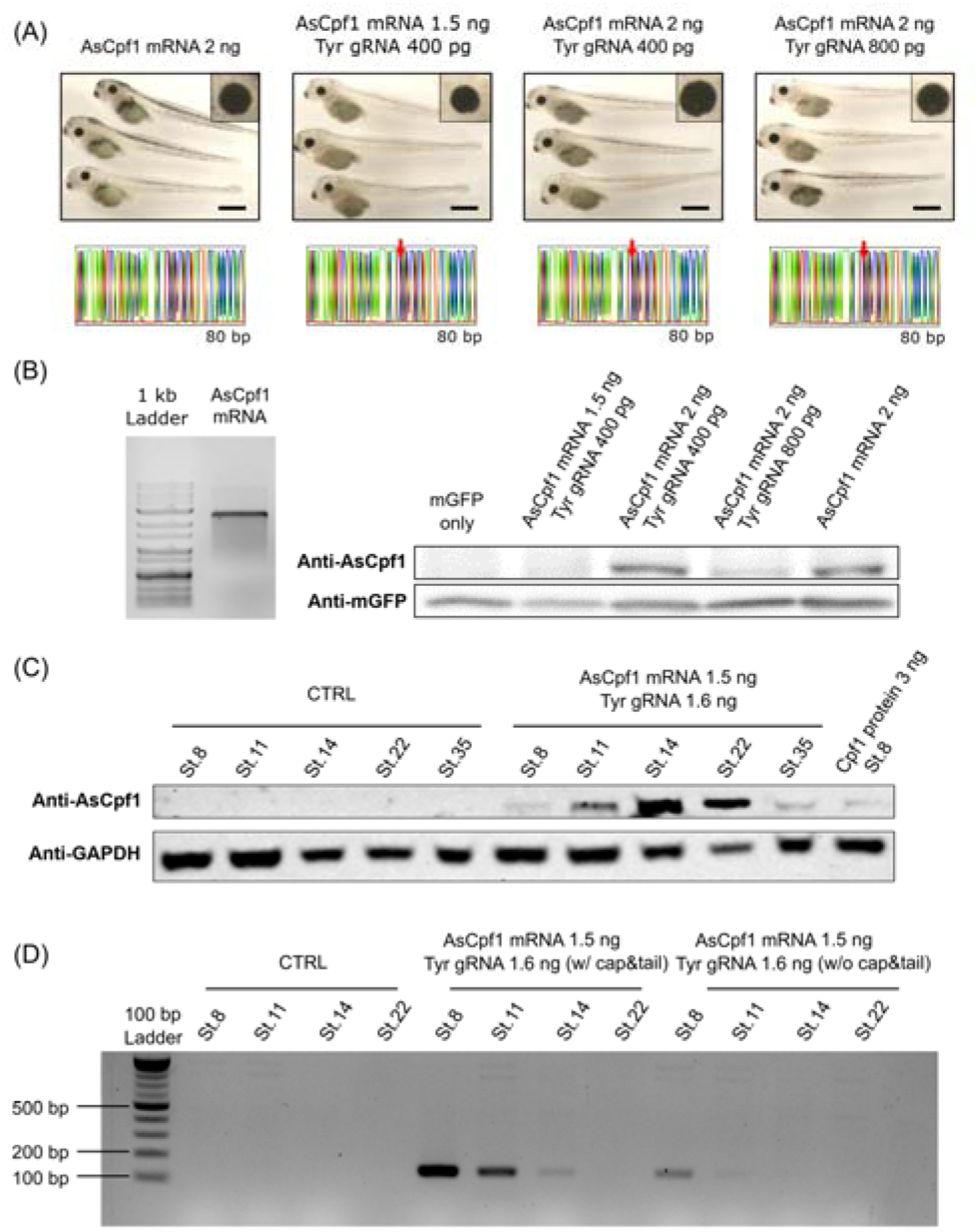
Microinjected AsCpf1-Ultra mRNA is non-functional. The efficacy of microinjected AsCpf1-Ultra mRNA against the *X. laevis* tyrosinase (*tyr*) gene was tested. **(A)** Wild-type tadpoles and all tadpoles injected with AsCpf1-Ultra mRNA and *tyr*-targeting gRNA at NF stage 40 showed complete pigmentation of the eye and skin, indicating that AsCpf1-Ultra does not affect the tyrosinase gene. Scale bar, 1 mm. Sanger sequencing of each embryo also did not show any noticeable genetic alteration induced by AsCpf1-Ultra. **(B)** AsCpf1-Ultra mRNA was synthesized by SP6 polymerase and confirmed by electrophoresis on an agarose gel. The *in vivo* protein expression level of AsCpf1-Ultra generated from the injected mRNA was measured by western blotting at NF stage 35-36 embryos. We compared the expression level with GFP proteins derived from co-injected mRNA. **(C)** We tested the expression of AsCpf1-Ultra in early developmental stages. It started to be expressed at Stage 8 and more proteins were translated at Stage 14 and Stage 22. However, the expression level is significantly reduced at Stage 35. **(D)** We quantified the availability of Cpf1 guide RNAs in IN4MER format (Table 3). Without any modification, it is only detected in early stage 8, then becomes not detectable after then. With a poly-A and capping modification like mRNA, it seems to be more stable, but still Stage 14 is the last stage for detection.

**Table 3.**
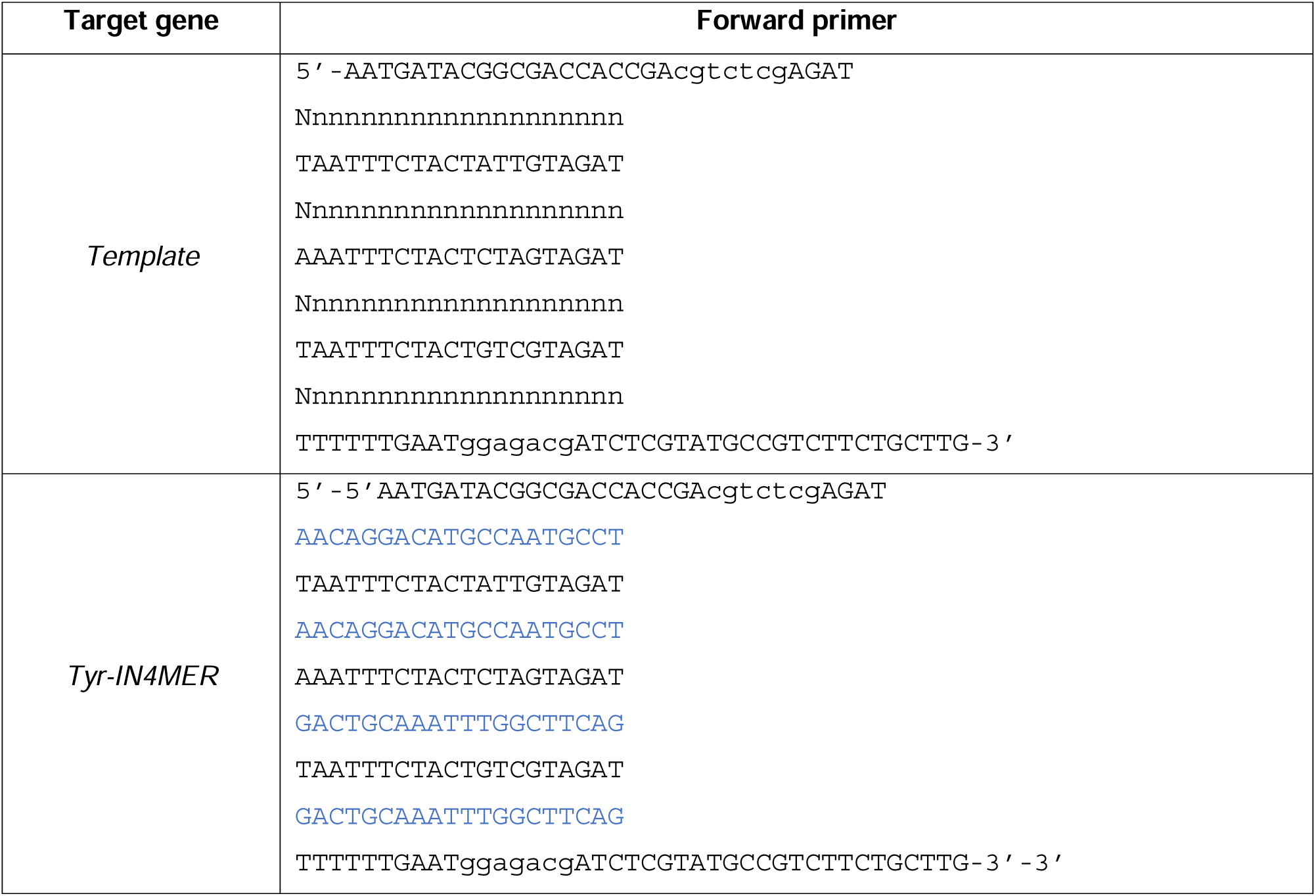
IN4MER template and sequence used in this study.

We also confirmed that there was no issue with the quality of the mRNA, and it was properly translated into protein by western blot from NF stage 35-36 embryos after two-cell stage injection (Fig. 4B). Furthermore, the same AsCpf1-Ultra mRNA induced DNA to break properly in a human cell line with the LNP delivery (data not shown). Therefore, we speculated that the stability of mRNA and guide RNAs may be different in *Xenopus* embryos. Protein translation takes time after mRNA injection, but if the guide RNAs are degraded rapidly, functional RNP cannot be formed.

Because the Cas12a guide RNA we used for the RNA experiment is 44 bp, it was too short to be measured with conventional quantitative RT-PCR. Recently, the IN4MER platform with a polycistronic guide RNA of 223 bp for Cas12a has been developed (Anvar et al. 2024). We designed IN4MER guide RNA with tyr-1 and tyr-2 guide RNAs (Table 3) and confirmed their functions through in vitro cleavage, similar to our previous experiment with the RNP, by incubating it with AsCpf1-Ultra proteins. As expected, both tyr-1 and tyr-2 targets were efficiently cleaved, confirming that the IN4MER platform works with AsCpf1-Ultra proteins. However, when we injected the IN4MER guide RNAs together with AsCpf1-Ultra mRNA into two-cell stage *X. laevis* embryos, we did not observe any phenotypic changes.

We further examined the degradation of the IN4MER guide RNA and AsCpf1-Ultra protein expression at earlier stages (Fig. 4C and Fig. 4D). Protein translation begins at NF stage 8, with more robust protein production occurring between NF stages 11 and 14. The amount of protein produced by 1.5 ng of AsCpf1-Ultra mRNA injection is much greater than what we can detect with 3 ng of protein injection, as shown in Fig. 4C, confirming that the amount of AsCpf1-Ultra protein is sufficient in *Xenopus* embryos after mRNA injection. However, when we measured the amount of the IN4MER guide RNAs with RT-PCR, we found that these guide RNAs are rapidly degraded after NF stage 8.

There is a report that poly-A tailing and 5’-capping, similar to mRNA transcripts, help stabilize guide RNA (Mu et al. 2019), so we tested the IN4MER guide RNAs with these modifications, which are the same as in AsCpf1-Ultra mRNA. Through *in vitro* cleavage assays, we confirmed that this modification does not affect the function of AsCpf1-Ultra (data not shown). The guide RNAs are much more stable at early NF stage 8, but they are also degraded by NF stage 14 and become undetectable after NF stage 22. From these results, we conclude that the rapid degradation of guide RNA before protein translation may be the reason for the lack of effective AsCpf1-Ultra activity when it was injected as mRNA.

## Discussion

CRISPR/Cas has become a widely used genome engineering tool in various animal models because of its convenience and high efficiency. Most of these applications are based on experiments in mammalian systems at 37°C, which seems optimal for the activity of the CRISPR/Cas system. However, this temperature is not suitable for all model organisms, particularly aquatic model organisms like African clawed frog *Xenopus* and zebrafish *Danio rerio*, which are usually raised at lower temperatures. This may limit the usability of CRISPR/Cas. Fortunately, SpCas9, the most widely used Cas enzyme in genome engineering, does not have a strong temperature dependency. Therefore, little attention has been paid to the temperature dependency of CRISPR applications. However, another Cas enzyme, Cas12a, exhibits activity at a narrow range of temperatures, and thus its use is limited in systems requiring lower temperatures without a heat-shock step to activate the Cas enzyme (Moreno-Mateos et al. 2017). This temperature dependency limitation was almost overcome by an engineered Cas12a enzyme with two amino acid substitutions (Zhang et al. 2021). However, its usability was not thoroughly investigated in aquatic model organisms that require activities at much lower temperatures than were tested in mammalian organisms. Here, we reported that engineered Cas12a enzymes (LbCpf1-Ultra and AsCpf1-Ultra) are highly active at the low temperature suitable for *Xenopus* cultivation (20–28°C). Furthermore, they outperformed SpCas9 in gene disruption and therefore could be a promising alternative to morpholinos for studying gene functions in aquatic model organisms.

Recently, Shi and colleagues comprehensively benchmarked the activities of various Cas9 and Cas12a enzymes in *X. tropicalis* (Shi et al. 2022). In addition to SaCas9 and KKH SaCas9, which are not widely used in this organism, they reported that an engineered Cas12a enzyme (EnGen-LbCpf1 from NEB) is highly effective at 23°C without heat shock. However, this enzyme was not as effective as other engineered Cas12a enzymes (LbCpf1-Ultra and AsCpf1-Ultra) in our *in vitro* digestion assay. One possible explanation for the low activity of EnGen-LbCpf1 in our study is that a higher concentration of this enzyme may be required to achieve similar activity, as in the case of SpCas9. Detailed information about how EnGen-LbCpf1 was engineered has not been disclosed; therefore, we cannot fully explain the difference between these engineered Cas12a enzymes. Further analysis with more Cas12a variants is necessary to elucidate the mechanism underlying the temperature dependency of this enzyme.

In contrast to the ribonucleoprotein complex of Cas12a and guide RNA, we observed that co-injecting Cas12a mRNA and guide RNA did not work in *Xenopus* embryos. Although this was previously reported in zebrafish (Moreno-Mateos et al. 2017), the molecular mechanism remains unclear. Because the guide RNA for Cas12a is too short to analyze with conventional Q-RT-PCR, investigating the stability of guide RNAs was challenging. Here, we used the polycistronic IN4MER guide RNA system (Anvar et al. 2024) and confirmed its rapid degradation in early-stage embryos. Since protein translation requires time after mRNA injection (until NF stage 8 to 11, roughly 6 to 16 hours after fertilization), it is crucial to retain a large amount of guide RNA in the cell. However, guide RNAs are rapidly degraded right after injection, making it difficult to maintain sufficient concentration to form the RNP complex after protein translation. Modifying guide RNA may help improve stability, but if the CRISPR RNP functions at a later stage, achieving gene disruption becomes more difficult compared to early-stage activity. Therefore, regardless of guide RNA stability, injecting the RNP complex will be a more efficient approach for gene disruption.

We observed phenotypes in F0 embryos without any selection; therefore, we expected stochastic and mosaic phenotypic penetrance, as reported with most SpCas9 crispants. Although we injected CRISPR/Cas before the first cell division in an embryo, it can cut the target sites after several rounds of cell division because the timing of its activities is not guaranteed. By performing quantitative phenotypic analysis, this study confirmed the surprisingly high gene disruption efficiency of engineered Cas12a in *Xenopus*. Previously, it was reported that the indel scars generated by DNA repair after *in vivo* DNA cleavage are much larger with Cas12a than with Cas9 (Shi et al. 2022). Furthermore, Cas12a makes four- or five-nucleotide staggered cuts in contrast with the blunt-ends generated by Cas9 (Zetsche et al. 2015) and therefore may cause more severe gene disruption compared with Cas9. We suggest that engineered Cas12a enzymes more effectively generate F0 phenotypes.

Cas12a is also reported to have unspecific single-strand DNA trans-cleavage activities (Chen et al. 2018), which could render this system toxic for genome engineering *in vivo*. An R-loop, a DNA-RNA hybrid related to transcriptional regulation, commonly forms during the cell cycle (Crossley et al. 2019) and is dynamically altered during development (Munden et al. 2022), so it could become a target of Cas12a trans-cleavage activity. Although R-loop formation during *Xenopus* development has not been fully elucidated, we observed no toxicity of engineered Cas12a during genome editing *in vivo.* Therefore, this effect may not occur, as reported in human cell lines (Kim et al. 2016; Kleinstiver et al. 2016) and even bacteria (Marino et al. 2022).

## Acknowledgements

This work was supported by the UNIST Research Fund (1.220050.01 and 1.230039.01 to TK; 1.250006.01 to SWC, TJP, and TK), by a National Research Foundation of Korea (NRF) grant funded by the Ministry of Education of the Korean government (RS-2018-NR031072 to TK), and by the Ministry of Science and ICT (2023R1A2C100627511 to TK, RS-2023-00213043 to SWC, and RS-2024-00354930 to TJP; RS-2024-00335111 to TJP and TK). It was also partially supported by a grant from the Institute for Basic Science (IBS-R022-D1 to SWC, TJP, and TK).

## Author Contributions

TK, TJP, and SWC conceived and designed the project. SY, SK, and JH mainly performed the experiments with the help of SS, JC, J-SS, SC, and HSL. SY, SK, JH, SWC, TJP, and TK analyzed the data and wrote the manuscript with the help of all authors.

